# Sperm regulates locomotor activity of *C. elegans* hermaphrodites

**DOI:** 10.1101/2023.05.02.539019

**Authors:** Satoshi Suo

**Affiliations:** Department of Pharmacology, Faculty of Medicine, Saitama Medical University, Saitama, Japan

## Abstract

Sex differences in sex-shared behavior are common across various species. During mating, males transfer sperm and seminal fluid to females, which can affect female behavior. Sperm can be stored in the female reproductive tract for extended periods of time. However, its role in regulating female behavior is poorly understood. In the androdioecious nematode *C. elegans*, hermaphrodites produce both egg cells and sperm, enabling them to self-fertilize or mate with males. Hermaphrodites exhibit less locomotor activity compared to males, indicating sex difference in behavioral regulation. In this study, sperm-deficient mutants were examined to investigate the role of sperm in the regulation of locomotor behavior. The results suggest that sperm plays a significant role in regulating sex-shared behavior in *C. elegans*, as sperm-deficient mutants exhibited increased locomotor activity. Additionally, females of closely related gonochoristic species, *C. remanei* and *C. brenneri*, exhibited reduced locomotor activity after mating. The regulation of locomotion by sperm may be an adaptive mechanism that enables sperm-less hermaphrodites to search for mates and allow females to cease their search for mates after mating.

## Introduction

There are sex differences in behavior, including mating behavior and shared behaviors such as locomotion[1–3]. Both internal and external stimuli are known to regulate these behaviors. Mating can influence the behavior of females, particularly in insects[4,5]. Males transfer sperm and seminal fluid to the female body through mating. Previous studies have indicated that seminal proteins may regulate the behavior of female flies, resulting in increased aggression and decreased receptivity to males after mating. Sperm can be stored in the female reproductive tract and these storages are called sperm reservoir. In wide-ranging animal species, including vertebrates and invertebrates, sperm reservoir has been shown to persist for extended periods of time[6,7]. However, the specific effects of the sperm reservoir on female behavior remain largely unknown.

*C. elegans* has two sexes: male and hermaphrodite[8]. C. elegans hermaphrodites are morphologically female, but produce both egg cells and sperm, allowing them to self-fertilize and also mate with males. Hermaphrodites produce sperm during their larval stage, store them in the spermatheca, and transition to producing eggs upon reaching adulthood. When males mate with hermaphrodites, their sperm swim to the spermatheca and at least partially displaces the hermaphrodite-derived sperm[9]. During fertilization, the male-derived sperm is preferred over hermaphrodite-derived sperm. Therefore, most of the progenies are born from cross-fertilization after mating.

*C. elegans* exhibits two distinct behavioral states, dwelling and roaming[10]. Dwelling animals move slowly and frequently change direction, whereas roaming animals exhibit higher locomotor activity and turn less frequently. There are sex differences in the regulation of these behavioral states, whereby hermaphrodites spend more time in the dwelling state, while males spend more time in the roaming state[11]. This allows males, which require mates to reproduce, to explore large areas to find hermaphrodites.

*C. elegans* hermaphrodites possessing self-sperm provides an opportunity to investigate the effect of stored sperm using sperm-deficient mutants without confounding effects of mating. In the present study, I examined the locomotion of sperm-deficient mutants to test whether sperm influences the regulation of behavioral states. The results showed that sperm-deficient mutants display elevated locomotor activity and increased roaming behavior, suggesting that the sperm reservoir plays a role in regulating sex-shared behavior in *C. elegans*. This study sheds new light on the potential contribution of internal sperm to the regulation of behavior.

## Materials and Methods

### Strains

Culturing of *C. elegans* was performed as previously described[8]. The strains used in this study are as follows: N2 wildtype (RRID:WB-STRAIN:N2), CB4088 *him-5(e1490)* (RRID:WB-STRAIN:CB4088), BA821 *spe-26(hc138)* (RRID:WB-STRAIN:BA821), BA-17 *fem-1(hc17)* (RRID:WB-STRAIN:BA17), CB4108 *fog-2(q71)* (RRID:WB-STRAIN:CB4108), CB3844 *fem-3(e2006)* (RRID:WB-STRAIN:CB3844), BA671 *spe-9(hc88)* (RRID:WB-STRAIN:BA671), EM464 *C. remanei* (RRID:WB-STRAIN:EM464), and CB5161 *C. brenneri* (RRID:WB-STRAIN:CB5161). CB4088 was used to obtain males for mating with *C. elegans* strains.

### Image acquisition and data analysis

The analyses of *C. elegans* locomotor behavior were performed as previously described[11] with some modifications. The assay plates were made of low peptone-NGM agar (0.25 g/L peptone, 3 g/L NaCl, 17 g/L Agar, 25 mM KPO4 (pH 6.0), and 5 mM MgSO4, 5 mM CaCl2). 10 μL of OP50 (RRID:WB-STRAIN:OP50) suspension was placed on the plates to create thin and small bacterial lawns.

L4 hermaphrodites (or females) were placed onto a 40 mm NGM plate[8] seeded with OP50 and were grown at 20°C for approximately 20 h. In the mating condition, hermaphrodites (or females) were cultured together with males of the same species. Subsequently, the animals were individually transferred to the assay plates 15 min prior to recording. The images of the assay plates were captured using a DMK series USB camera (Imaging Source, Bremen, Germany) with a resolution of 2,000 × 1,944 pixels, at one frame per sec for 25 min, using the gstreamer software with Raspberry Pi 3 (Raspberry Pi Foundation, Cambridge, United Kingdom).

The position of the animals within the bacterial lawn was obtained using a custom-written code in Python and the average speed and angular speed were determined. To analyze behavioral states, the average speed and angular speed of 10-sec intervals were determined and plotted[11]. Data points located above a predefined line were classified as roaming and those below were categorized as dwelling, similarly as described in a previously study[12].

### Sperm staining

Males were stained with MitoTracker Red CMXRos (ThermoFisher, Waltham, MA) as previously described[13]. L4 males were picked onto MitoTracker containing plates, prepared by placing 100 μL of 10 μM Mitotracker on OP50-seeded NGM plates. After growing 16 h at 20°C, adult males were transferred to new plates without Mitotracker and mated with adult hermaphrodites (or females) for 6 h. The hermaphrodites were then mounted on slides and their images were taken using a fluorescence microscope (BX53, Olympus, Tokyo, Japan).

### Statistical analysis

Numbers of animals tested for each experiment are shown in the figure legends. Statistical analysis was performed using Python. The Wilcoxon rank-sum test was used to determine the p values. The Bonferroni correction was used for comparisons of more than two groups. For comparisons of behavioral states, the percent of time spent in roaming state was compared between strains and conditions. The differences were considered statistically significant with ^*^p < 0.05, ^**^p < 0.01, ^***^p < 0.001. For the box plots, the boxes depict the lower and upper quartile values, and the central lines represent the medians. The whiskers are the most extreme values within 1.5 times the interquartile range and the dots represent outliers.

## Results

### Sperm-less animals exhibit increased locomotor activity

In this study, I investigated the locomotor behavior of *C. elegans* animals cultured on plates containing food bacteria. We recorded the behavior of individually placed animals and determined the average speed and average angular speed of 10-second intervals. Consistent with previous studies[11,12,14], wildtype hermaphrodite animals exhibited low speed and high angular speed. High angular speed indicates frequent turning of animals. Data points were plotted (Fig 1 A, B), with many points located in the low-speed and high-angle region, and fewer points located in the high-speed and low-angle region. Similar to a previous study[12], we separated the behavioral states. Points with high speed and low angular speed were categorized as the roaming state, while points with low speed and high angular speed were categorized as the dwelling state. Our findings confirmed that wildtype animals spent most of their time in the dwelling state.

**Figure 1.**
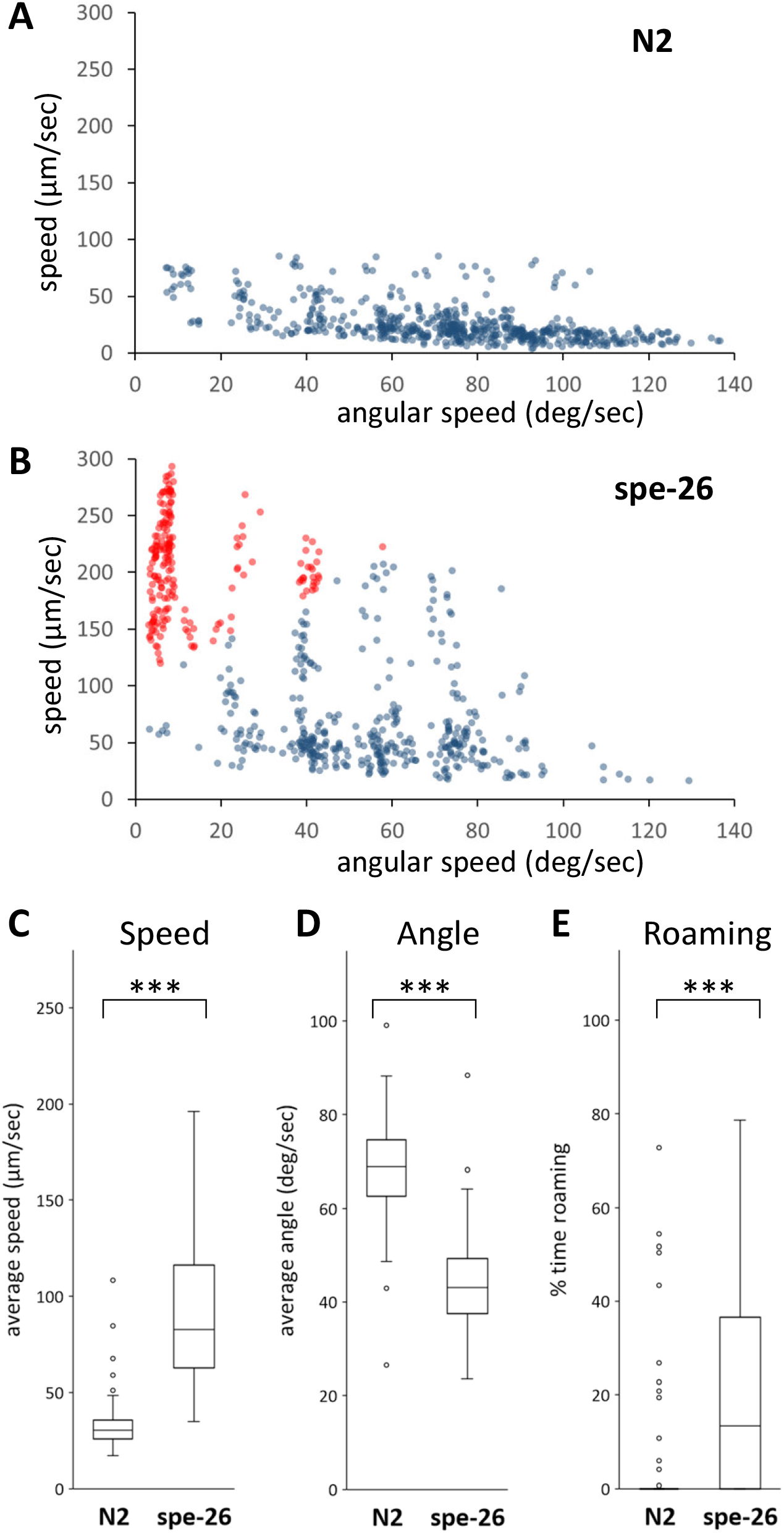
Locomotor behavior of N2 and *spe-26* hermaphrodites. Scatter plots of average speed and average angular speed in 10-sec intervals for 16 each N2 hermaphrodites (A) and sperm-deficient *spe-26* mutant hermaphrodites (B). Data points were categorized into roaming (red) and dwelling (blue). Average speed (C), angular speed (D), and the percentage of time spent in the roaming state (E) were determined. Numbers of the animals tested: N2, 75; *spe-26*, 79.

Hermaphrodites of the sperm-deficient *spe-26* mutant[15] were tested to investigate the effect of sperm on locomotor behavior. We recorded and plotted the data points for the *spe-26* hermaphrodites and found that *spe-26* hermaphrodites exhibited a greater number of points with high-speed and low-angle, when compared to their wildtype counterparts. The *spe-26* hermaphrodites exhibited increased speed, decreased angular speed, and increased time spent in the roaming state, when compared to wildtype hermaphrodites (Fig 1C-E). These results suggest that the presence of sperm suppresses roaming behavior and the loss of sperm results in an increase in locomotor activity.

In addition to the *spe-26* mutants, I also examined other sperm-less mutants. The *fem-1, fem-3*, and *fog-1* mutants have defects in sex-determination genes that are required to promote spermatogenesis and therefore produce oocytes instead of sperm[16–18]. In contrast, spermatogenesis appears to complete in *spe-9* mutants, but the *spe-9* gene is required for sperm function [19]. I examined the locomotor behavior of the hermaphrodites of these mutants and found that all of them had significantly increased speed and decreased angular speed compared to wildtype hermaphrodites. Furthermore, the roaming behavior was also increased in *fem-1, fog-2*, and *spe-9* mutants. As several mutants that cause loss of sperm function in different manners each showed increased locomotor activity, the results further support the notion that sperm suppresses locomotion in hermaphrodites.

### Mating suppresses increased locomotion of sperm-deficient animals

The effect of adding sperm to hermaphrodites on locomotion was investigated to determine whether sperm can suppress increased locomotion in sperm-less mutants. Male-derived sperm moves to the spermatheca after mating and can be stored for an extended period[9]. To confirm the presence of male-derived sperm in hermaphrodites, MitoTracker-stained males were mated with wildtype and *spe-26* hermaphrodites, and fluorescent cells, most likely sperm, were observed in both wildtype and *spe-26* hermaphrodites (Fig 3). For the behavioral analysis, the movements were recorded from both wildtype and *spe-26* hermaphrodites that had either mated or not mated with males. There were less data points with high speed and low angular speed in mated *spe-26* mutants compared to unmated *spe-26* mutants (Fig 4A,B). *spe-26* mutants exhibited increased speed, decreased angle, and increased roaming compared to wildtype hermaphrodites. However, these locomotor changes were reduced by mating (Fig 4C-E). In contrast, in wildtype hermaphrodites, mating did not significantly alter their speed, angle, and roaming. These results demonstrate that mating suppresses the increased locomotion of sperm-less animals, further suggesting that stored sperm inside hermaphrodites suppresses locomotion.

**Figure 2.**
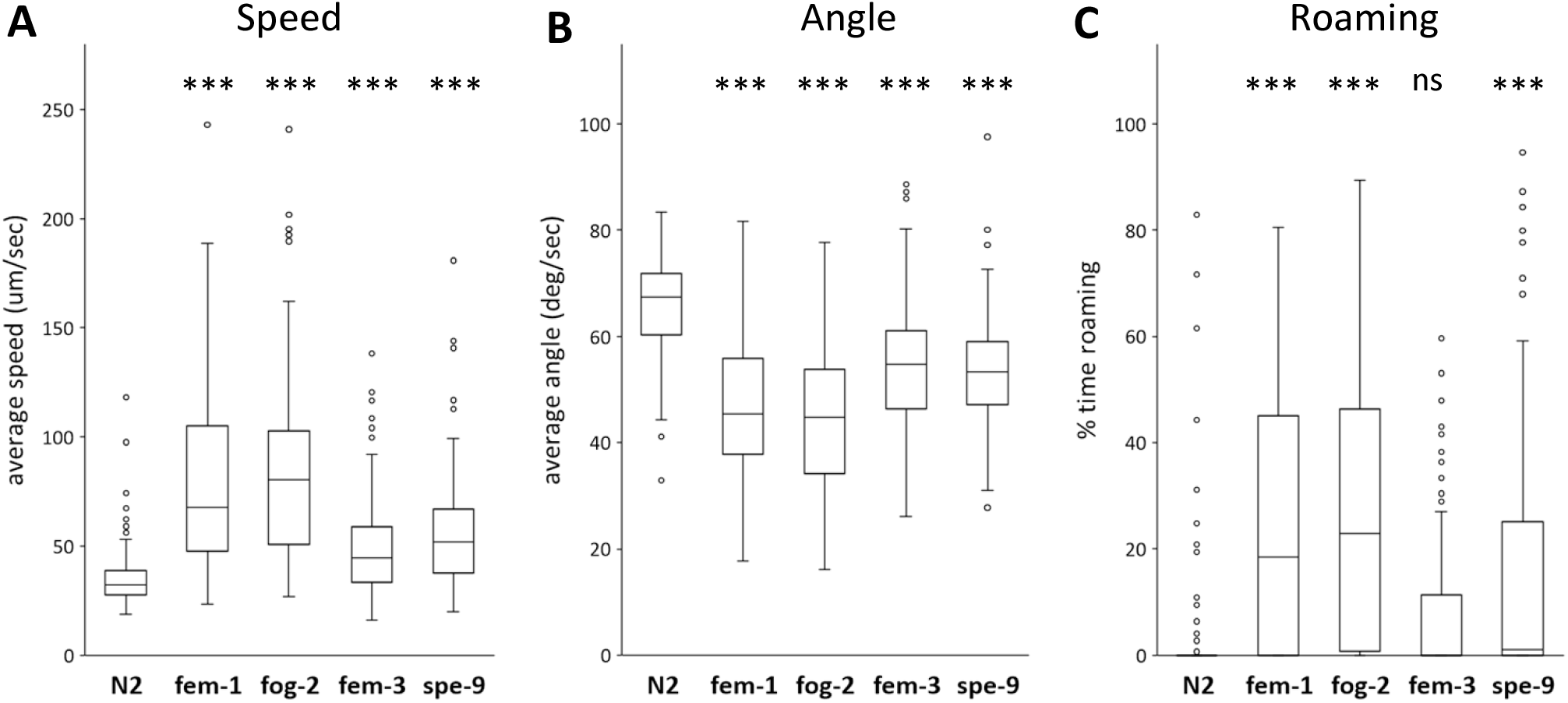
Locomotor behavior of sperm-deficient mutants. Average speed (A), angular speed (B), and the percentage of time spent in the roaming state (C) of the *fem-1, fog-2, fem-3*, and *spe-9* mutants were determined. Numbers of the animals tested: N2, 91; *fem-1*, 93; *fog-2*, 91; *fem-3*, 92; *spe-9*, 92.

**Figure 3.**
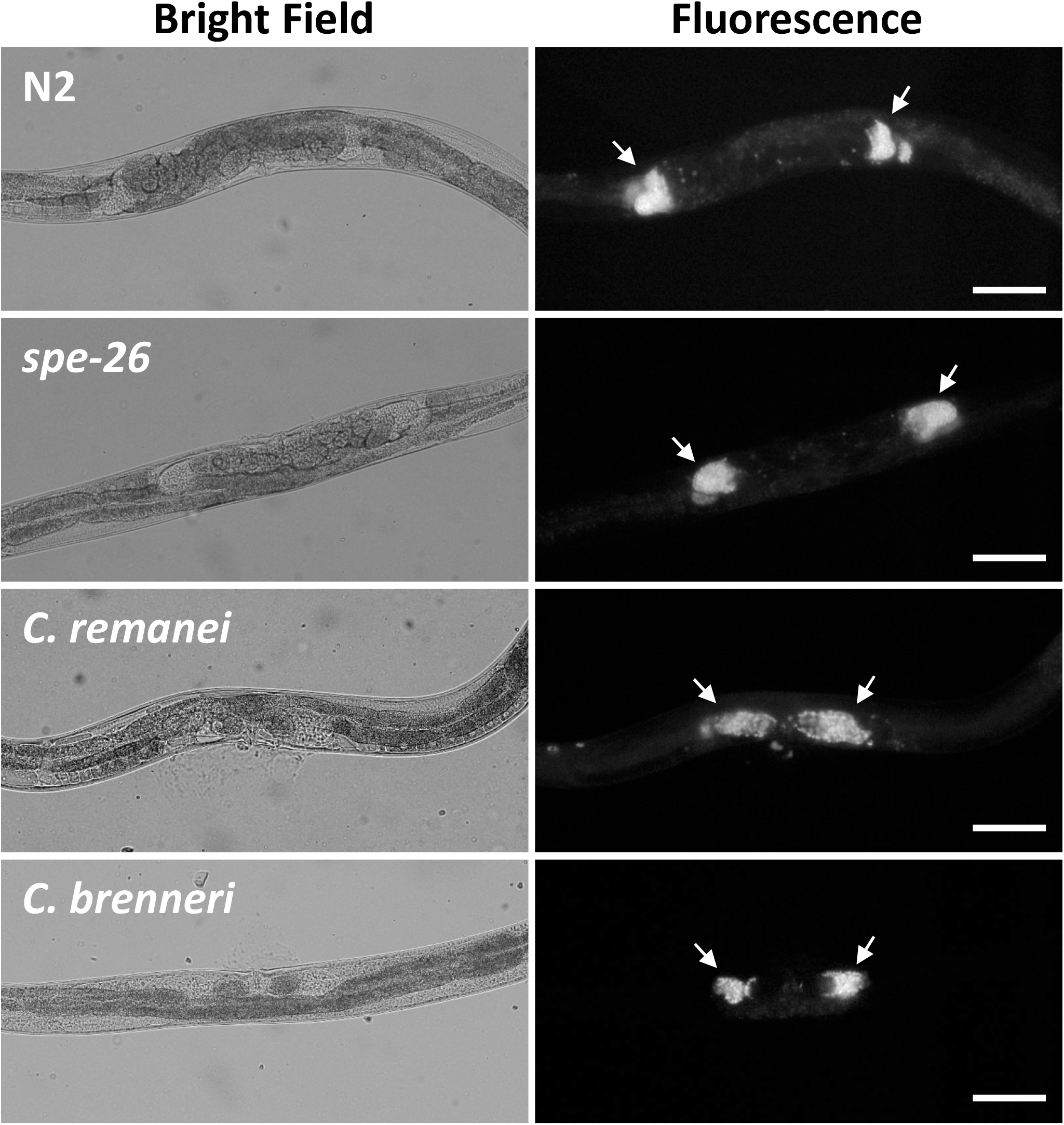
Transfer of sperm. N2 and *spe-26 C. elegans* hermaphrodites, and *C. remanei* and *C. brenneri* females were mated with corresponding males stained with MitoTracker, and the bright field and fluorescence images were obtained. Arrows indicate possible sperm fluorescence accumulating at the location of spermatheca. Scale bar = 10 μm.

**Figure 4.**
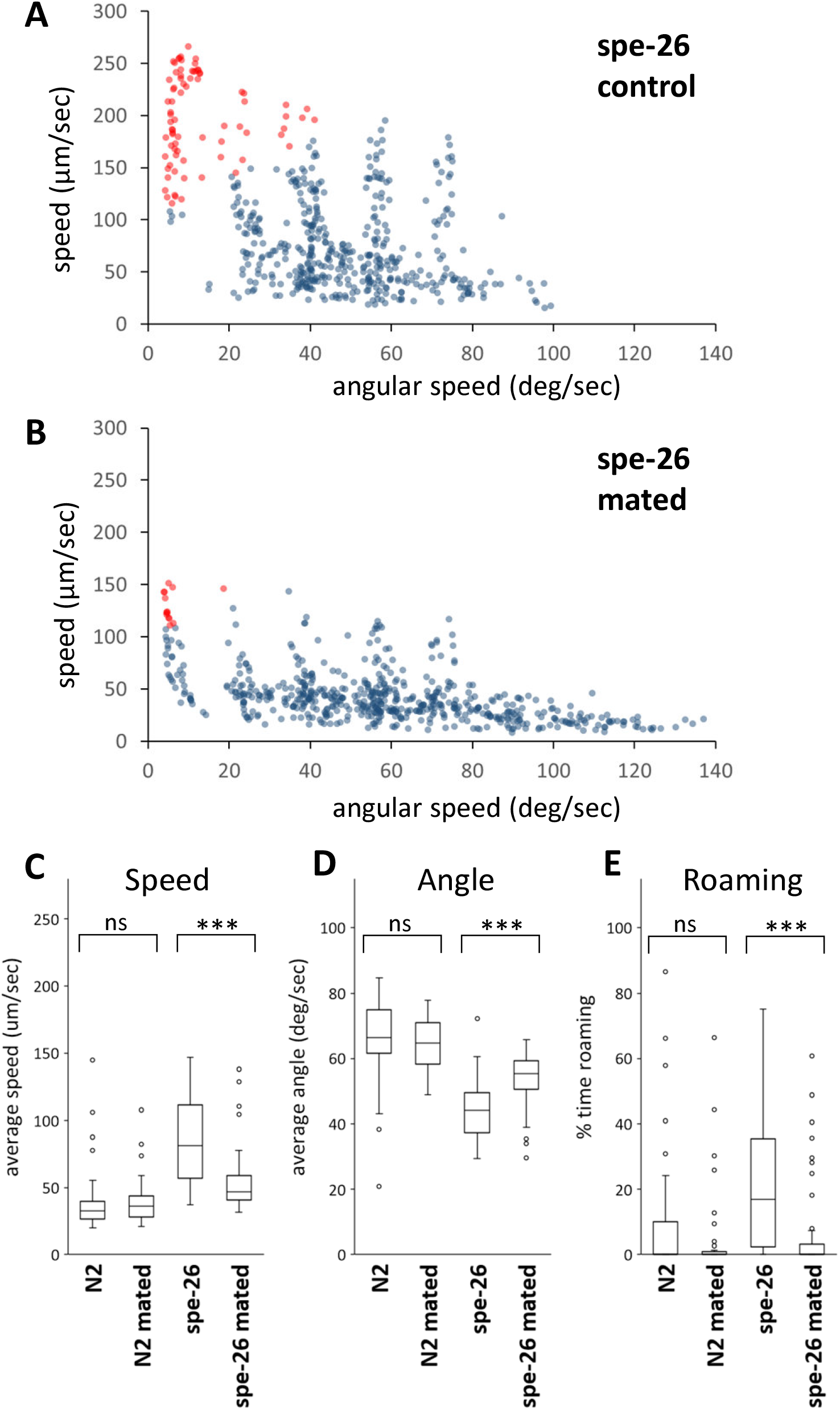
Locomotor behavior of N2 and *spe-26* hermaphrodites after mating. Scatter plots of average speed and average angular speed in 10-sec intervals for 16 each of control *spe-26* hermaphrodites (A) and *spe-26* hermaphrodites mated with males (B). Data points were categorized into roaming (red) and dwelling (blue). Average speed (C), angular speed (D), and the percentage of time spent in the roaming state (E) were determined. Numbers of the animals tested: N2 control, 47; N2 mated, 45; *spe-26* control, 61; *spe-26* mated, 64.

### Mating suppresses locomotion in females of other *Caenorhabditis* species

*C. elegans* is an androdioecious species, which has males and hermaphrodites, whereas other closely related species, such as *C. remanei* and *C. brenneri*, are gonochoristic and have male and female sexes. I also investigated whether mating regulates locomotion in these gonochoristic species. First, dye-stained males were mated with females, and it was confirmed that male-derived sperm is stored in the female reproductive tracts (Fig 3). Behavioral analyses revealed that females of these gonochoristic species exhibit a higher locomotor rate than *C. elegans* hermaphrodites. Furthermore, there were a decrease in speed, an increase in angle, and an increase in roaming after mating with males in *C. remanei* and *C. brenneri* females, similar to what was observed in the sperm-deficient *C. elegans* mutants (Fig 5). These results suggest that the behavior of female nematodes changes after mating, possibly due to sperm transfer.

**Figure 5.**
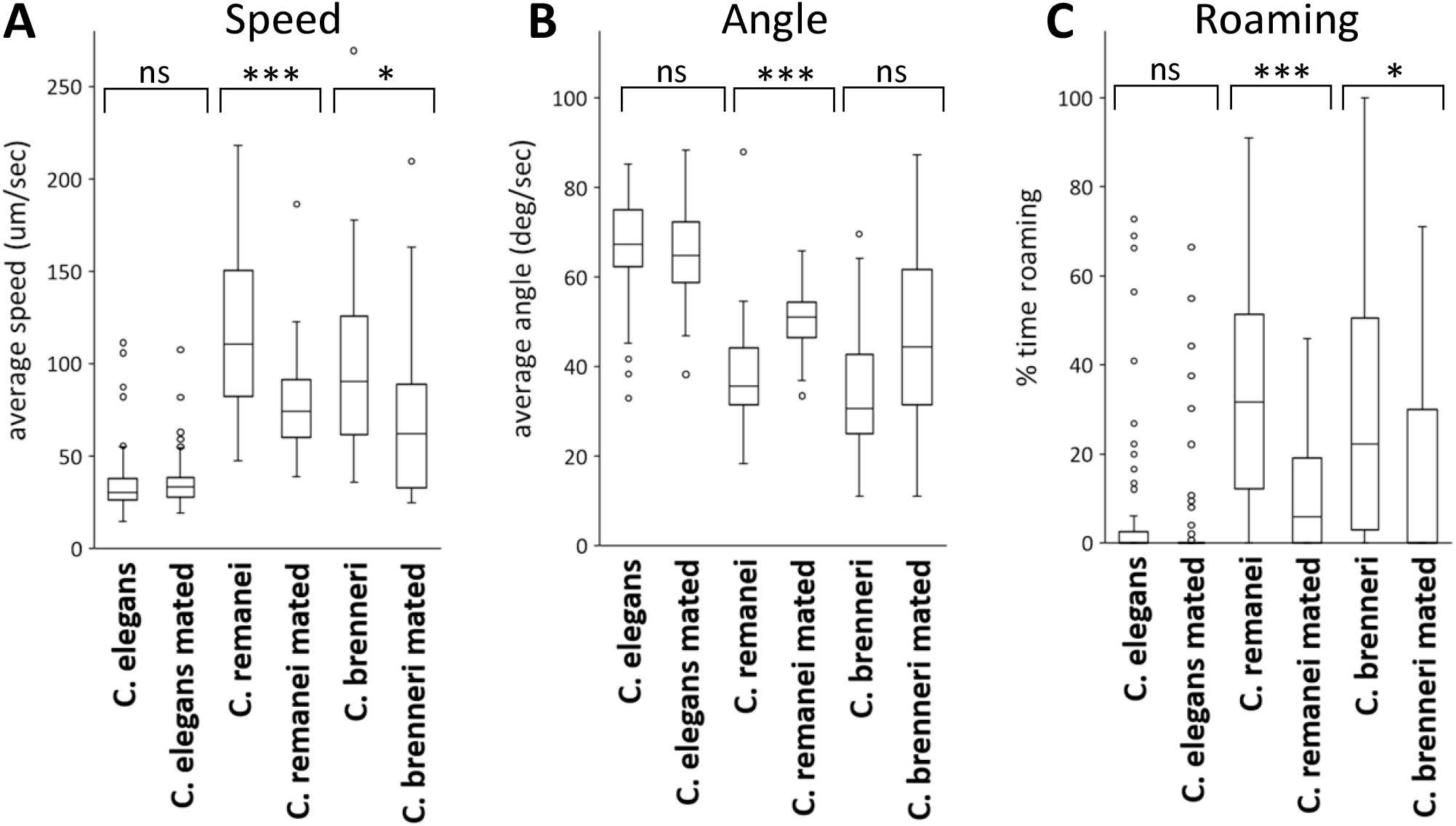
Locomotor behavior of *Caenorhabditis* species after mating. Average speed (A), angular speed (B), and the percentage of time spent in the roaming state (C) of *C. elegans* hermaphrodites, *C. remanei* females, and *C. brenneri* females were determined. For mated, animals were mated with corresponding males prior to recording. The average speed of unmated animals was higher in *C. remanei* (p < 0.01) and *C. brenneri* (p < 0.01), when compared to *C. elegans*. Numbers of the animals tested: *C. elegans* control, 76; *C. elegans* mated, 76; *C. remanei* control, 61; *C. remanei* mated, 74; *C. brenneri* control, 39; *C. brenneri* mated, 46.

## Discussion

This study demonstrated that the hermaphrodites of sperm-deficient mutants exhibit increased locomotor activity compared to wildtype hermaphrodites. The consistent increase in locomotor activity across multiple mutants with defects in sperm development supports the idea that sperm plays a role in suppressing locomotor activity in hermaphrodites. The *fem-1, fem-3*, and *fog-2* mutants have defects in germ-line sex determination and produce oocytes instead of sperm[16–18]. The *spe-26* mutants have a defect in the early stages of sperm development and produce aberrant spermatocytes that do not progress to spermatids[15]. In contrast, *spe-9* mutants produce spermatozoa with wild-type morphology, but these sperm are not functional and cannot fertilize oocytes[19]. These results suggest that the presence of mature functional sperm is necessary for suppressing locomotor activity in adult hermaphrodites.

The effect of mating on hermaphrodite’s behavior was also examined. The results showed that sperm-less *spe-26* mutants exhibit a reduction in locomotor activity after mating with males, which results in the transfer of male-derived sperm in the spermatheca where hermaphrodite’s self-sperm is normally stored. This further supports the idea that the presence of sperm suppresses locomotor activity in *C. elegans*. However, these findings do not rule out the possibility that other factors also play a role in regulating locomotor activity. Mating can trigger modifications in behavior and physiology through the transfer of seminal fluid[4,5]. In addition, the chemical and physical perception of males may also play a role in this process. Therefore, it is possible that the observed decrease in locomotor activity in this study is not solely mediated by sperm but could be a result of combination of various factors.

In our previous study, we observed that *C. elegans* males exhibit higher levels of locomotor activity and spend more time in the roaming state than hermaphrodites[11]. This sex difference in locomotion may be adaptive, as males require mates to reproduce and therefore benefit from exploring larger areas to find potential mates. In contrast, hermaphrodites are capable of self-fertilization and may conserve energy by remaining in areas with ample food. The present study showed that the loss of sperm results in increased locomotor activity of hermaphrodites. Sperm-less hermaphrodites, which effectively are females, require males to reproduce. Therefore, this behavioral change may be adaptive, as it makes the sperm-less hermaphrodites to actively search for mates.

There are male-female species of nematodes closely related to *C. elegans*. Notably, the females of two such strains, *C. remanei* and *C. brenneri*, were more active than *C. elegans* hermaphrodites. Furthermore, a decrease in locomotor activity and roaming behavior was observed in mated females, who received sperm from males and effectively became hermaphrodite. This behavioral regulation would allow females to conserve energy and remain near available food sources after mating, when they no longer need to search for mates.

Sperm status affects global gene expression in *C. elegans*[20] and alters aspects of hermaphrodite physiology in *C. elegans*. Furthermore, sperm facilitates the maturation of oocytes[21], increases cold tolerance[22], protect from mating-induced death[23,24] and reduces attractiveness to males[25]. This study expands on these findings and demonstrate that sperm plays a role in regulating locomotor behavior. The mechanism by which hermaphrodites detect sperm to regulate their behavior is currently unknown. One possibility is that sperm itself or factors released from sperm is detected by hermaphrodite cells. The major sperm protein released from sperm is detected by receptor proteins on oocytes and gonad cells, which in turn regulates oocyte maturation[21]. Another possibility is that fertilized eggs are somehow detected by hermaphrodites. Further research is required to determine the molecular mechanisms underlying the suppression of locomotor activity by signaling from sperm to the nervous system.

In summary, the results suggest that sperm plays an important role in regulating locomotor activity and behavioral states in nematodes. Loss of sperm in hermaphrodites leads to increased activity, potentially allowing effective females to seek mates. These findings highlight the adaptive significance of sperm-regulation of sex-shared behavior.

## Acknowledgements

The nematode strains were provided by the Caenorhabditis Genetics Center, which is funded by NIH Office of Research Infrastructure Programs (P40 OD010440).

## References

1. Mowrey WR, Portman DS. Sex differences in behavioral decision-making and the modulation of shared neural circuits. Biol Sex Differ. 2012;3: 8. doi:10.1186/2042-6410-3-8

2. McCarthy MM. Multifaceted origins of sex differences in the brain. Phil Trans R Soc B. 2016;371: 20150106. doi:10.1098/rstb.2015.0106

3. Auer TO, Benton R. Sexual circuitry in Drosophila. Current Opinion in Neurobiology. 2016;38: 18–26. doi:10.1016/j.conb.2016.01.004

4. Kubli E, Bopp D. Sexual Behavior: How Sex Peptide Flips the Postmating Switch of Female Flies. Current Biology. 2012;22: R520–R522. doi:10.1016/j.cub.2012.04.058

5. Bath E, Bowden S, Peters C, Reddy A, Tobias JA, Easton-Calabria E, et al. Sperm and sex peptide stimulate aggression in female Drosophila. Nat Ecol Evol. 2017;1: 0154. doi:10.1038/s41559-017-0154

6. Suarez SS. Mammalian sperm interactions with the female reproductive tract. Cell Tissue Res. 2016;363: 185–194. doi:10.1007/s00441-015-2244-2

7. Pascini TV, Martins GF. The insect spermatheca: an overview. Zoology (Jena). 2017;121: 56– 71. doi:10.1016/j.zool.2016.12.001

8. Brenner S. The genetics of Caenorhabditis elegans. Genetics. 1974;77: 71–94.

9. LaMunyon CW, Ward S. Larger sperm outcompete smaller sperm in the nematode Caenorhabditis elegans. Proc Biol Sci. 1998;265: 1997–2002.

10. Fujiwara M, Sengupta P, McIntire SL. Regulation of Body Size and Behavioral State of C. elegans by Sensory Perception and the EGL-4 cGMP-Dependent Protein Kinase. Neuron. 2002;36: 1091–1102. doi:10.1016/S0896-6273(02)01093-0

11. Suo S, Harada K, Matsuda S, Kyo K, Wang M, Maruyama K, et al. Sexually Dimorphic Regulation of Behavioral States by Dopamine in Caenorhabditis elegans. J Neurosci. 2019;39: 4668–4683. doi:10.1523/JNEUROSCI.2985-18.2019

12. Ben Arous J, Laffont S, Chatenay D. Molecular and Sensory Basis of a Food Related Two-State Behavior in C. elegans. PLoS ONE. 2009;4: e7584. doi:10.1371/journal.pone.0007584

13. Hu M, Legg S, Miller MA. Measuring Sperm Guidance and Motility within the Caenorhabditis elegans Hermaphrodite Reproductive Tract. J Vis Exp. 2019. doi:10.3791/59783

14. Stern S, Kirst C, Bargmann CI. Neuromodulatory Control of Long-Term Behavioral Patterns and Individuality across Development. Cell. 2017;171: 1649-1662.e10. doi:10.1016/j.cell.2017.10.041

15. Varkey JP, Muhlrad PJ, Minniti AN, Do B, Ward S. The Caenorhabditis elegans spe-26 gene is necessary to form spermatids and encodes a protein similar to the actin-associated proteins kelch and scruin. Genes Dev. 1995;9: 1074–1086.

16. Schedl T, Kimble J. fog-2, a germ-line-specific sex determination gene required for hermaphrodite spermatogenesis in Caenorhabditis elegans. Genetics. 1988;119: 43–61. doi:10.1093/genetics/119.1.43

17. Ahringer J, Kimble J. Control of the sperm-oocyte switch in Caenorhabditis elegans hermaphrodites by the fem-3 3’ untranslated region. Nature. 1991;349: 346–348. doi:10.1038/349346a0

18. Spence AM, Coulson A, Hodgkin J. The product of fem-1, a nematode sex-determining gene, contains a motif found in cell cycle control proteins and receptors for cell-cell interactions. Cell. 1990;60: 981–990. doi:10.1016/0092-8674(90)90346-g

19. Singson A, Mercer KB, L’Hernault SW. The C. elegans spe-9 gene encodes a sperm transmembrane protein that contains EGF-like repeats and is required for fertilization. Cell. 1998;93: 71–79. doi:10.1016/s0092-8674(00)81147-2

20. Angeles-Albores D, Leighton DHW, Tsou T, Khaw TH, Antoshechkin I, Sternberg PW. The Caenorhabditis elegans Female-Like State: Decoupling the Transcriptomic Effects of Aging and Sperm Status. G3 (Bethesda). 2017;7: 2969–2977. doi:10.1534/g3.117.300080

21. Han SM, Cottee PA, Miller MA. Sperm and oocyte communication mechanisms controlling C. elegans fertility. Developmental Dynamics. 2010;239: 1265–1281. doi:10.1002/dvdy.22202

22. Sonoda S, Ohta A, Maruo A, Ujisawa T, Kuhara A. Sperm Affects Head Sensory Neuron in Temperature Tolerance of Caenorhabditis elegans. Cell Reports. 2016;16: 56–65. doi:10.1016/j.celrep.2016.05.078

23. Shi C, Booth LN, Murphy CT. Insulin-like peptides and the mTOR-TFEB pathway protect Caenorhabditis elegans hermaphrodites from mating-induced death. Elife. 2019;8: e46413. doi:10.7554/eLife.46413

24. Booth LN, Maures TJ, Yeo RW, Tantilert C, Brunet A. Self-sperm induce resistance to the detrimental effects of sexual encounters with males in hermaphroditic nematodes. Elife. 2019;8: e46418. doi:10.7554/eLife.46418

25. Morsci NS, Haas LA, Barr MM. Sperm Status Regulates Sexual Attraction in Caenorhabditis elegans. Genetics. 2011;189: 1341–1346. doi:10.1534/genetics.111.133603

